# Conclusive Identification of Senescent T Cells Reveals Their Abundance in Aging Humans

**DOI:** 10.1101/2020.06.17.157826

**Authors:** Ricardo Iván Martínez-Zamudio, Hannah K. Dewald, Themistoklis Vasilopoulos, Lisa Gittens-Williams, Patricia Fitzgerald-Bocarsly, Utz Herbig

## Abstract

Aging leads to a progressive functional decline of the immune system, which renders the elderly increasingly susceptible to disease and infection. The degree to which immune cell senescence contributes to this functional decline, however, remains unclear since methods to accurately identify and isolate senescent immune cells are missing. By measuring senescence-associated ß-galactosidase activity, a hallmark of senescent cells, we demonstrate here that healthy humans develop senescent T lymphocytes in peripheral blood with advancing age. Particularly senescent CD8+ T cells increased in abundance with age, ranging from 30% of the total CD8+ T cell population in donors in their 20s and reaching levels of 64% in donors in their 60s. Senescent CD8+ T cell populations displayed features of telomere dysfunction-induced senescence as well as p16-mediated senescence, developed in various T cell differentiation states and established gene expression signatures consistent with the senescence state observed in other cell types. On the basis of our results we propose that cellular senescence of T lymphocytes is a major contributing factor to the observed decline of immune cell function with advancing age and that immune cell senescence, therefore, plays a significant role in the increased susceptibility of the elderly to age-associated diseases and infection.

## INTRODUCTION

Cellular senescence is a stable proliferative arrest that is encountered by mammalian cells in response to a variety of signals and stresses (*1*). In mammals, this response functions to suppress cancer development and plays also important roles during tissue repair, wound healing and embryonic development (*2*). Although senescent cells (SCs) can be cleared from tissue by adaptive and innate immune responses under these circumstances, not all SCs are cleared and consequently accumulate progressively in various tissues during aging (*3*). In mouse models, this age-associated accumulation of SCs has a substantial negative impact on fitness and health, as it shortens lifespan and contributes to the development of numerous age-associated diseases (*4, 5*). While it is currently unclear why some SCs evade immune cell clearance and accumulate in tissues, it is possible that immune cells themselves progressively become senescent with age, thereby increasingly weakening immune responses that would otherwise clear SCs from aged tissue and suppress infections from viruses and other pathogens (*3*). This hypothesis, however, has proven challenging to test, primarily due to the lack of a suitable marker that can identify senescent mammalian immune cells accurately and efficiently (*6, 7*).

In mammalian cells, at the least two pathways activate the senescence program. One pathway, mediated by the tumor suppressor p53 and cyclin-dependent kinase (CDK) inhibitor p21, primarily becomes activated in response to a persistent DNA damage response (DDR), such as the one caused due to telomere dysfunction. In normal somatic human cells, or cells that lack detectable telomerase activity, dysfunctional telomeres are generated not only due to repeated cell division cycles that cause progressive telomere erosion, but also as a consequence of genotoxic stresses that cause double stranded DNA breaks (DSBs) in telomeric repeats (*8, 9*). A second senescence pathway is activated due to upregulation of the CDK4/6 inhibitor p16^INK4a^, which results in a stable pRb-dependent cell cycle arrest. While genotoxic stresses *can* activate this senescence pathway in certain cell types, p16^INK4a^ is also upregulated in the absence of telomere dysfunction or a persistent DDR (*10, 11*). Although the molecular triggers of this DDR independent senescence response are still largely unclear, p16^INK4a^ mediated-senescence clearly has important physiological consequences, as it significantly contributes to aging and the development of age-related disorders in mammals (*5*).

Despite differences in pathway activation, SCs share a number of features. One characteristic that is common to all SCs is that they secrete numerous NF-*k*B and p38-MAPK regulated pro-inflammatory cytokines and other molecules, collectively called the senescence associated secretory phenotype or SASP (*12*). Although the SASP differs in composition depending on cell type, senescence-inducing signal, and time elapsed following senescence induction, a primary function of the SASP is to generate a pro-inflammatory environment that stimulates an immune response(*3*). Another feature common to SCs is that they up-regulate certain heterochromatin proteins, such as macroH2A, leading to a stable repression of cell proliferation genes (*13, 14*). In addition, SCs develop a greater abundance of lysosomal content and a reduction in lysosomal pH (*15*), resulting in increased expression of lysosomal ß-galactosidase. This hallmark of SCs in particular allows their detection in cultures and in tissue, regardless of the senescence-inducing signal or senescence pathway activated (*16*).

Our current understanding of cellular senescence stems primarily from studies conducted using mammalian fibroblast cultures. Senescence pathways in other cell types, including those of circulating peripheral blood mononuclear cells (PBMCs) CD4+ T cells, CD8+ T cells, monocytes, B cells natural killer (NK) cells, and plasmacytoid dendritic cells (pDC’s), are still incompletely understood (*17*). Although subsets of PBMCs, such as cytotoxic CD8+ T cells undergo replicative senescence in culture, whether and to what degree they do so also *in vivo* remains unclear (*7, 18*). A primary reason for this uncertainty is that specific markers used to identify senescent CD8+ T cells in the past (*19, 20*), such as a loss of the cell surface receptors CD28 and CD27 and a gain of expression of CD45RA, CD57, TIGIT and/or KLRG1 do not accurately characterize all T cells that have permanently lost the ability to proliferate due to acquisition of macromolecular damage, upregulation of cyclin-dependent kinase (CDK) inhibitors, and development of senescence associated ß-Galactosidase (SA-ßGal), criteria that define the state of cellular senescence (*16*). In fact, CD8+ T cells that have lost expression of CD28 and/or that display varying levels of CD45RA, CD57, TIGIT or KLRG1 maintain the ability to proliferate following appropriate stimulation, which is incompatible with a classical senescence response (*18, 19*).

Here, we describe an accurate and efficient method to quantify, isolate, and characterize live senescent immune cells from peripheral blood of human donors. We reveal the identity of PBMC subsets that increasingly undergo cellular senescence in healthy humans with age, uncover causes for cellular senescence in circulating CD8+ T cells, and characterize the pathways activated in senescent CD8+ T cells at levels that may provide insights into therapeutic opportunities to modulate T cell senescence in disease, infection, and advanced age.

## RESULTS

### A method to isolate live SCs for subsequent analysis

One feature that is common to SCs is that they develop increased expression levels SA-ßGal (*16*). This hallmark of SCs allows their detection using chromogenic (X-Gal) (*21*) or fluorogenic (FDG) (*22*) ßGal substrates, albeit with limitations due to poor cell permeability or lack of cellular retention, respectively. To mitigate these limitations, we tested a cell permeable and self-immobilizing fluorogenic SA-ßGal substrate (fSA-ßGal) for its ability to label live SCs for prolonged periods, so that they can be accurately analyzed, quantified, and isolated by flow cytometry. As anticipated, incubation with fSA-ßGal caused GM21 fibroblasts in oncogene-induced senescence (OIS), either alone or mixed at a 3:1 ratio with non-senescent GM21 fibroblasts, to develop significantly increased fluorescence signal intensities compared to proliferating non-senescent control fibroblasts (Fig. S1A). Live cells in OIS could be accurately sorted and isolated from non-SCs by FACS, allowing us to conduct cell proliferation assays, immunofluorescence analysis, and gene expression profiling. We show that FACS isolated cells with high SA-ßGal activity (fSA-ßGal high) were significantly impaired in their ability to proliferate, suppressed expression of cyclin A, and displayed increased expression levels of senescence genes *p21, p16^INK4a^, IL1B*, and *IL8* compared to cells with low activity (fSA-ßGal low; Fig. S1B-E). Similar results were obtained when comparing etoposide-treated fibroblasts undergoing DNA damage-induced senescence with proliferating non-senescent fibroblasts (Fig. S1F-H). Collectively, these results demonstrate that fSA-ßGal is specific to labeling live human SCs and efficient for isolating them from mixed cell populations for subsequent analysis.

### Subsets of human PBMCs increasingly develop high SA-ßGal activity with advancing age

To determine whether subsets of human peripheral blood mononuclear cells (PBMCs) undergo cellular senescence as a function of age in circulation, we collected and analyzed blood from healthy human donors in two age groups: 1) “young”, which included donors between the ages 23 and 30 years (average age 25 years; 20s), and 2) “old” which included donors between the ages 57 and 67 years (average age 64 years; 60s), around the age the human immune system is found to exhibit age-associated deficits. Young and old donors were always recruited in pairs, allowing us to process and analyze freshly isolated PBMCs from donor pairs in parallel (Fig. 1A and Fig. S2A). Cord blood, together with blood from healthy donors in their 20s, was also collected and analyzed for the presence of SCs. Surprisingly, age-associated increases of mean fluorescence intensities (MFIs) of SA-ßGal signals were observed in all subsets analyzed for some donor pairs, including T lymphocytes, plasmacytoid dendritic cells (pDC’s), natural killer (NK) cells, monocytes, and B cells (Fig. 1B-C and Fig. S2B). Our data therefore suggest that all subsets of human PBMCs can undergo cellular senescence *in vivo*. Most consistent age-associated increases in mean fluorescence intensities (MFIs), however, were observed only for T cells (Fig. S2B-C). In order to determine which PBMC subsets increasingly develop high SA-ßGal activity with advancing age, we quantified the percentages of cells that display increased fSA-ßGal signal intensities by taking advantage of the fact that young donors, and often also old donors, had distinct populations of cells with lower and higher fSA-ßGal fluorescence. This allowed us to set gates at intersections where the two populations met, designating the population with lower signal intensity as “fSA-ßGal low” and the population with higher signal intensity as “fSA-ßGal high” (Fig. 1B). Quantitative analysis of fSA-ßGal high cells revealed statistically significant age-associated increases in human T lymphocytes, particularly in CD8+ T cells. For this subset, we discovered a striking age-associated increase of cells with high SA-ßGal activity, ranging from 30 ± 3 % in blood from donors in their 20s to 64 ± 4 % in donors in their 60s (Fig. 2C; p<0.0001). CD4+ T cells also displayed a significant age associated increase of cells with high fSA-ßGal activity, albeit to a lesser degree compared to CD8+ T cells (15 ± 3 % in young and 31 ± 6 % in old donors, p=0.02). No gender differences in the percent of fSA-ßGal high cells were observed in either T cell population (not shown). Not surprisingly, percentages of cells with high SA-ßGal activity were the lowest in cord blood, including in T lymphocytes, which contained fewer that 5% of this SA-ßGal expressing population (Fig. 1C and Fig. S2D-E). Our data therefore demonstrate that an increasing fraction of T lymphocytes, in particular CD8+ T cells, develop high SA-ßGal activity with age.

**Fig. 1.**
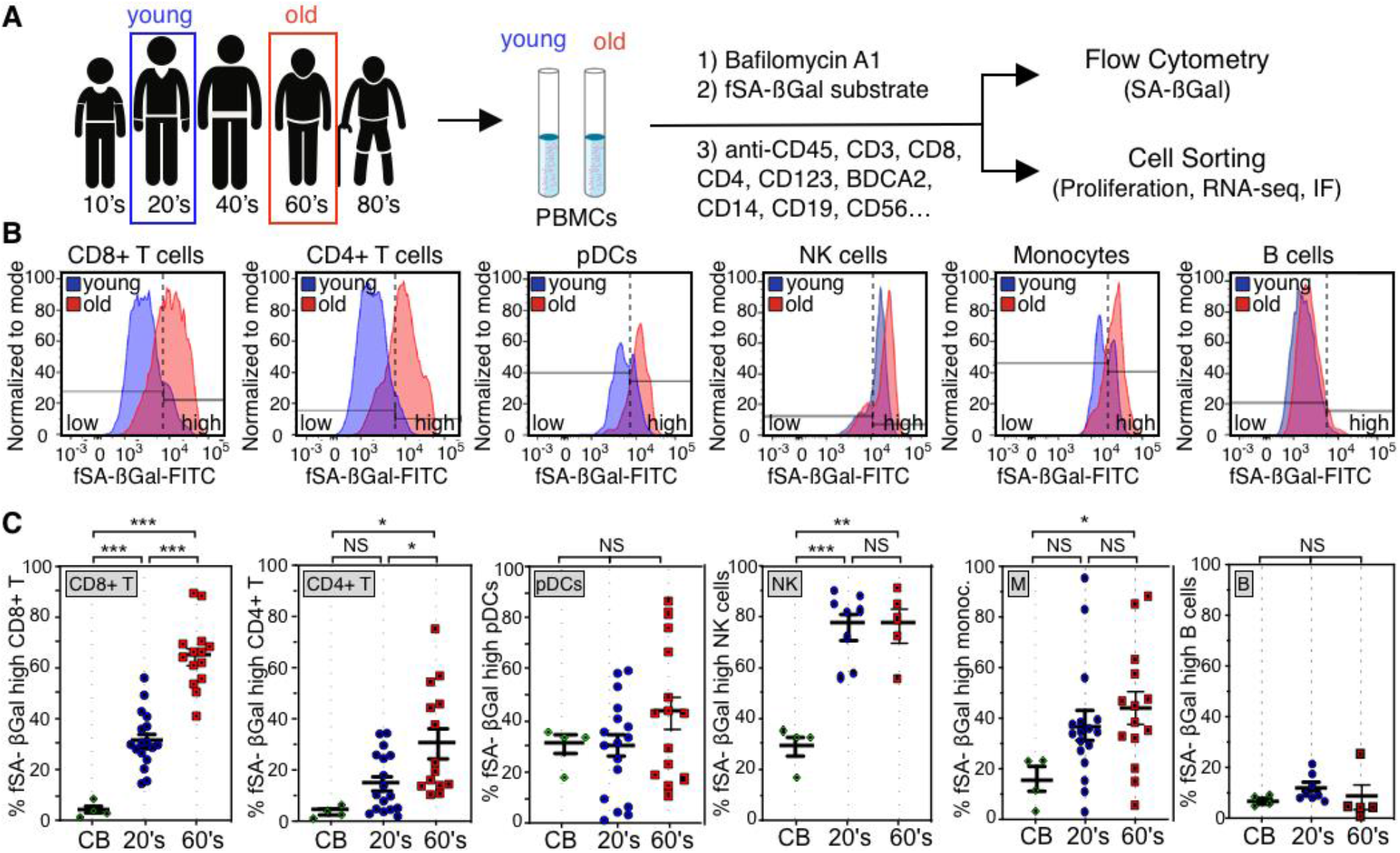
Humans display increased percentages of T lymphocytes with high fSA-ßGal signal intensities in advanced age. (**A**) Experimental strategy to quantify, isolate, and characterize senescent subsets of PBMCs from donors in their 20s (young) and 60s (old). IF: immunofluorescence analysis. (**B**) Representative fSA-ßGal intensity profiles and gates used to quantify fSA-ßGal high cells for indicated PBMC subsets from a young (blue) and old (red) donor. (**C**) Quantification of the percentages of fSA-ßGal high cells in cord blood (CB), young (blue) and old (red) donors for indicated PBMC subsets. Whiskers indicate mean +/- S.E.M and are indicated for each subset. Statistical significance was determined by an unpaired, two-tailed Student’s t test.*** p<0.0001; ** p = 0.0005; * p < 0.05; NS: not significant.

### CD8+ T cells with high levels of SA-ßGal activity display features of telomere dysfunction-induced senescence (TDIS) and p16^INK4a^-mediated senescence

To test whether CD8+T cells with high SA-ßGal activity are senescent, we sorted and collected them by FACS based on low, intermediate, or high fSA-ßGal signal intensities (Fig. 2A) and measured cell proliferation, senescence gene expression, and development of features specific to SCs. Significantly, the ability of CD8+T cells to proliferate following activation was inversely proportional to fSA-ßGal signal intensities. While 91 ± 1% of cells with low SA-ßGal activity were able to undergo more than one cell division during a 5 day period of stimulation, the fractions of proliferating cells with intermediate and high fSA-ßGal signal intensities were reduced to 71% ± 6% and 33 ± 6%, respectively, under the same conditions (Fig. 2B). No significant differences in proliferation capacities were observed between T cells collected from young or old donors at respective fSA-ßGal signal intensities (Fig. 3A). Gene expression analysis by RT-qPCR demonstrated significantly increased expression levels of senescence genes p16^INK4a^ and p21, in cells with high SA-ßGal activity compared to cells with intermediate or low activities (Fig. 2C), which is consistent with a senescence response. Levels of IL6 mRNA, however, decreased with SA-ßGal activity, which is unlike senescent human fibroblasts (*12*). In addition, immunofluorescence analysis revealed that protein levels of p16^INK4a^ and the senescence marker macroH2A were the lowest in CD8+ T cells with low SA-ßGal activity and increased proportionally in cells with intermediate and high SA-ßGal activities (Fig. 2D and Fig. S3B-C). Similarly, cells that displayed multiple foci of 53BP1, a DDR factor that localizes to sites of DSBs, were less frequently detected in cells with low SA-ßGal activity, compared to cells with intermediate or high activities, irrespective of donor age (Fig. 2D-E and Fig. S3D-F). Approximately 40% of DDR foci analyzed were localized at telomeric repeats, revealing that a substantial fraction of DSBs in CD8+ T cells are caused as a result of telomere dysfunction (Fig. 2F and Fig. S3G). Overall, the mean number of telomere dysfunction-induced DNA damage foci (TIF) per cell increased proportionally with increasing SA-ßGal activity, irrespective of donor age, demonstrating a direct correlation between the senescence state and presence of dysfunctional telomeres in CD8+ T cells (Fig. 2G and Fig. S3G-H). Our data therefore demonstrate that humans increasingly develop senescent CD8+ T cells with features of TDIS and p16^INK4a^-mediated senescence with age.

**Fig. 2.**
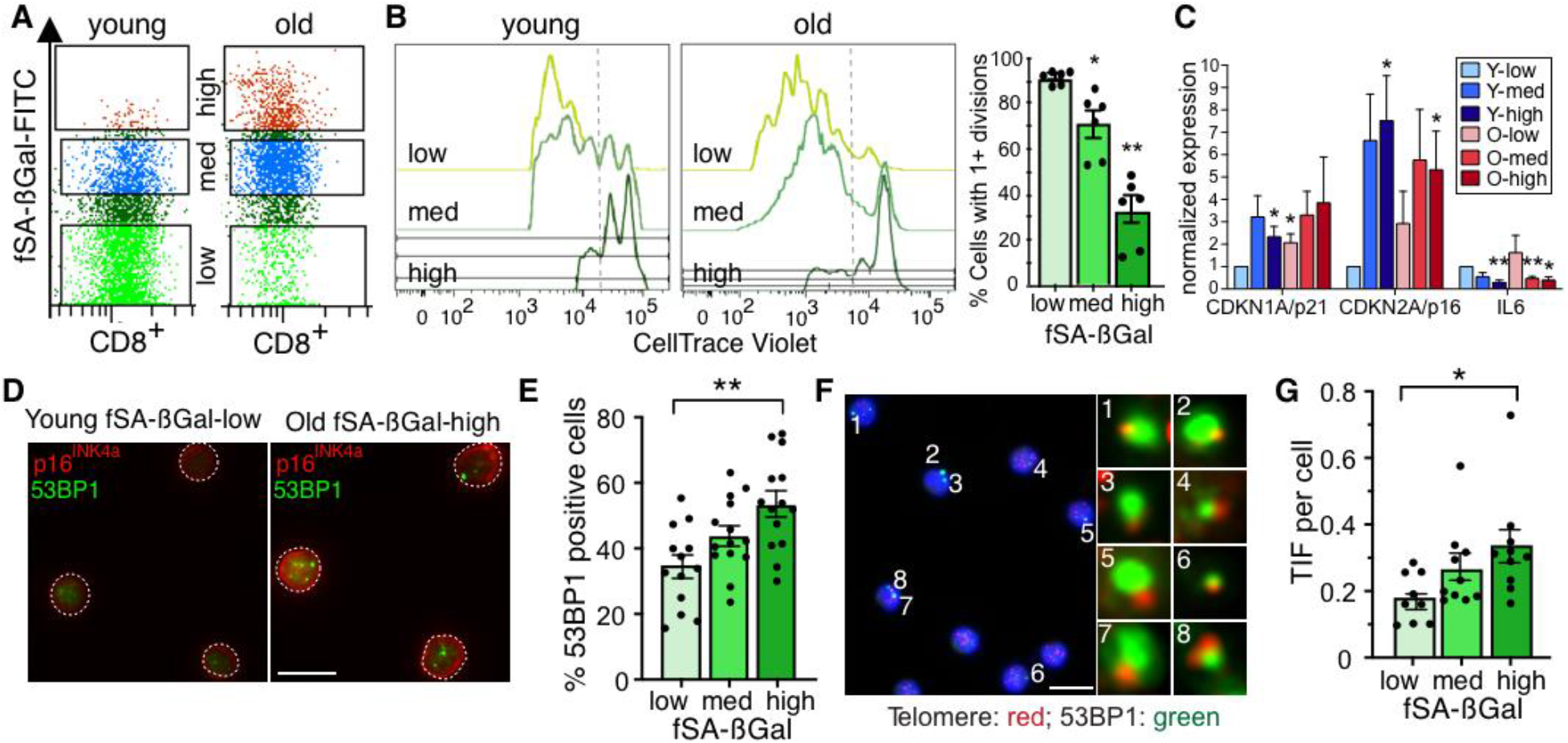
CD8+ T cells with high fSA-ßGal signal intensities are senescent. (**A**) Dot plot illustrating gates used to sort and isolate CD8+ T cells by FACS from young and old donors, as indicated, based on low, inter(med)iate, and high fSA-ßGal signal intensities. (**B**) Representative Cell Trace Violet histograms of CD8+ T cells, sorted as in (a), following anti-CD3 and anti-CD28 stimulation for 5 days from a young and old donor. Bar graph, quantification of more than one cell divisions of CD8+ T cells from 6 healthy donors (3 young, 3 old) following polyclonal stimulation for 5 days. n = 3; * p = 0.0002; ** p < 0.0001. (**C**) RT-qPCR expression profiles for indicated senescence-associated genes in CD8+ T cells at indicated fSA-ßGal levels in young (blue) and old donors (red). Average and S.E.M. of five (CDKN1A and CDKN2A) and three (IL6) independent experiments. Statistical significance was determined by a two-tailed unpaired t-test, with Y-low as the reference value. * p < 0.05; ** p< 0.001. (**D**) Immunofluorescence analysis of sorted CD8+ T cells from a young and old donor, as indicated, using antibodies against 53BP1 (green) and p16^INK4a^ (red). Outline of cell nuclei is indicated with white border. Scale bar: 10 □m (**E**) Quantification of the percentage of CD8+ T cells positive for 53BP1 foci at each fSA-ßGal level in young and old donors combined. Young: n = 8, Old: n = 8. Whiskers depict mean +/- S.E.M. Statistical significance was determined by a one-way ANOVA. ** p = 0.001. (**F**) Sorted CD8+ T cells with high fSA-ßGal signal intensities were simultaneously immunostained using antibodies against 53BP1 (green) and analyzed by FISH to detect telomeres (red). Blue: DAPI. Enlarged versions of the numbered DNA damage foci showing colocalization with telomeres are shown in the right micrographs. Scale bar: 10 □m (**G**) Quantification of mean TIF per cell in sorted CD8+ T cells, as indicated, in young and old donors combined. Young: n = 5, Old: n = 5. Whiskers depict mean +/- S.E.M. Statistical significance was calculated by a one-way ANOVA. * P = 0.0280.

### Senescent CD8+ T cells develop a unique gene expression signature

To characterize the phenotype of senescent CD8+ T cells in greater detail, we performed RNA-sequencing on CD8+ T cells with high and low SA-ßGal activities isolated from 3 young and 3 old healthy human donors. Significantly, principal component analysis (PCA) and hierarchical clustering revealed that the variability between the 12 transcriptomes was largely determined by SA-ßGal activity, not by donor age (Fig. 3A and Fig. S4A). A total of 4,149 genes were differentially expressed between senescent and non-senescent CD8+ T cells, corresponding to 2,362 up-regulated and 1,787 down-regulated genes with a minimal fold-change of 1.2. Weighted correlation network analysis (WGCNA)(*23*) generated two gene modules, yellow and blue, displaying high co-expression interconnectivity (Fig. S4B-D). Consistent with the PCA, differentially expressed genes within each module partitioned based on whether cells displayed high or low SA-ßGal activity and not based on donor age (Fig. 3B). Functional overrepresentation profiling enriched for a highly diverse array of biological processes consistent with a senescence response in the yellow module, including ‘p53 pathway, ‘Inflammatory response’, and ‘TGFß signaling’, many which overlapped with pathways also activated in senescent human fibroblasts (*24, 25*) (Fig. S4E-F). In contrast, the blue module was overall less diverse, enriching for pathways that included ‘IL2- STAT5 signaling’, ‘IL6-JAK-STAT3 signaling’ and ‘MYC targets’, which are consistent with cell proliferation and normal CD8+ T cell function (*26*) (Fig. 3C). Significantly, 71% of the 4,149 differentially expressed genes (DEGs) of senescent CD8+ T cells were also differentially expressed in senescent human fibroblasts, demonstrating a substantial overlap in senescent gene regulation between the two cell types (Fig. 3D). Interestingly, cellular senescence of CD8+ T cells resembled a state of prolonged or deep senescence, as DEGs and pathways activated showed greater overlap between senescent CD8+ T cells and fibroblasts that had undergone replicative senescence for extended periods (4 months), rather than for a shorter period (2 months; Fig. 3D and Fig. S4F) (*24*). Our data therefore raise the possibility that that senescent CD8+ T cells persist in circulation for months or longer.

**Fig. 3.**
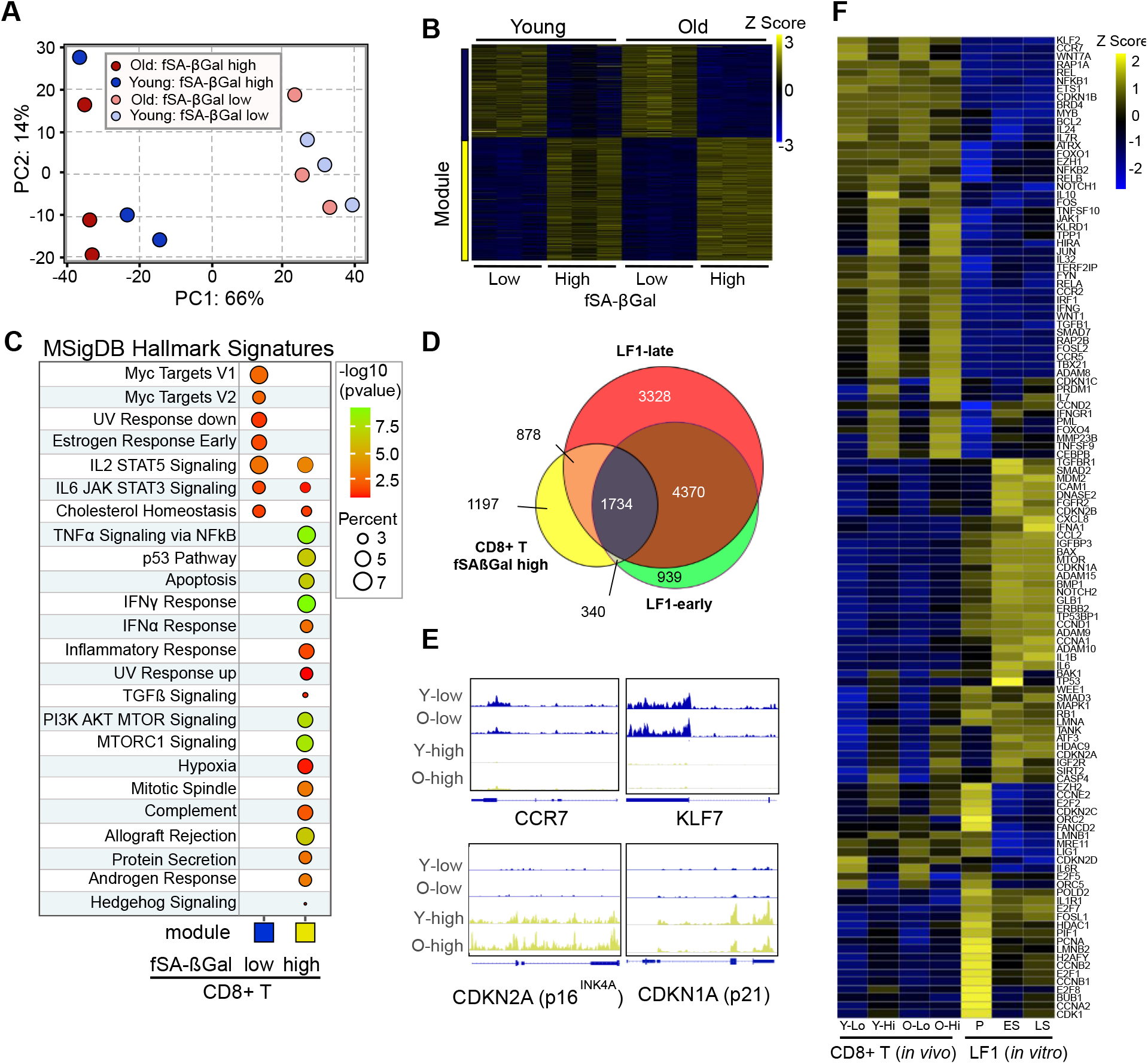
CD8+ T cells with high fSA-ßGal signal intensities develop a gene expression signature that is characteristic of senescent human cells. (**A**) Principal component analysis on transcriptomes from SA-ßGal high and low CD8 cells isolated from 3 old and 3 young donors. (**B**) Expression heatmap of the DEGs within each module across fSA-ßGal-low and -high CD8+ T cells. Each column represents an individual donor. Data are represented as Z-scores. (**C**) Functional over-representation map depicting Molecular Signatures Database (MSigDB) hallmark gene sets associated to each transcriptomic cluster. Circles are color-coded according to the FDR-corrected p-value based on the hypergeometric distribution test. (**D**) Venn diagram portraying the intersections and disjunctive unions of differentially expressed genes in CD8+ T cells with high fSA-ßGal activity (yellow) and human lung fibroblasts (LF1) that had remained in replicative senescence for 2 months (LF1-early; green) and 4 months (LF1-late; red). (**E**) Representative genome browser visualizations of normalized reads at indicated gene loci in CD8+ T cells sorted based on low and high fSA-ßGal signal intensities from a representative young (y) and old (o) donor, as indicated, using RNA-seq. (**F**) Expression heatmap of a selection of senescence-associated genes in CD8+ T cells isolated from peripheral blood and sorted based on low (Lo) and high (Hi) fSA-ßGal signal intensities from young (Y) and old (O) donors and *in vitro* cultured LF1 fibroblasts in proliferation (P), early (ES; 2 months) and late senescence (LS; 4 months) as in (**D**) Each column represents the average expression of 3 independent donors (CD8+) and 3 independent experiments (LF1).

A more detailed comparison of the expression of a subset of senescence-associated genes revealed commonalities and cell type-specific senescence gene expression profiles. Classical senescence-associated genes, such as products of the *CDKN1* and *CDKN2* loci were expressed in both cell types, albeit at different magnitudes (Fig. 3E-F). Expression of SASP genes, however, was more cell type-specific. For instance, interleukins (*IL*)*1A, 1B* and *6*, were highly expressed in senescent fibroblasts but not in senescent CD8+ T cells. In contrast, expression levels of *TNF, TGFB1* and *IFNG* were high in senescent CD8+ T cells, but not in senescent fibroblasts (Fig. S4G). Overall, our data therefore demonstrate that senescent human CD8+ T cells develop a unique senescence gene expression signature *in vivo*, yet a signature that overlaps substantially with that of other *in vitro* grown human cell types.

### CD8+ T cells increasingly undergo cellular senescence with advancing age, irrespective of their differentiation state

CD8+ T cell populations can be divided into distinct subsets based on their functions, and the expression of cell surface receptors CCR7 and CD45RA, among others (*27*). Subsets include naïve (T_N_: CCR7+/CD45RA+), central memory (T_CM_: CCR7+/CD45RA-), effector memory (T_EM_: CCR7-/CD45RA-), and effector (T_EMRA_: CCR7-/CD45RA+) T cells. Although TEMRA cells have been shown to display some features of cellular senescence, such as increased DNA damage, as well as reduced proliferative capacity, telomere lengths, and telomerase activity (*28, 29*), it remains unclear whether they can be defined as senescent in the classical sense, particularly as T_EMRA_ cells retain function and their growth arrest is reversible (*6, 20*). In line with previous studies (*19*), we observed that the fraction of CD8+ T_EM_, and T_EMRA_ cells were significantly more abundant in old donors compared to young donors or cord blood, while the fraction of TN cells wasdramatically decreased with age (Fig. 4A and Fig. S5A-B). Surprisingly, cells with high SA-ßGal activity were detected, albeit in different proportions, in all differentiation states, regardless of donor age (Fig. 4A and Fig. S5B). While young donors developed SCs primarily in T_EM_ subsets, old donors displayed the greatest fraction of SCs in both, the T_EM_ and T_EMRA_ subsets (Fig. 4A-C and Fig. S5C). However, a significant percentage of T_EMRA_ cells (89 %, 38 %, and 20 % in cord blood, young, and old donors, respectively) did not display increased SA-ßGal activity, demonstrating that this differentiation state is comprised of both senescent and non-SCs (Fig. S5C). Importantly, a fraction of T_N_ cells also displayed high SA-ßGal activity, particularly in old donors where 17.3 +/- 3 % T_N_ cells were found in the fSA-ßGal high population. This demonstrates that cellular senescence affects CD8+ T cells in all differentiation states, including in naïve T cells (Fig. 4A-C and Fig. S5C).

**Fig. 4.**
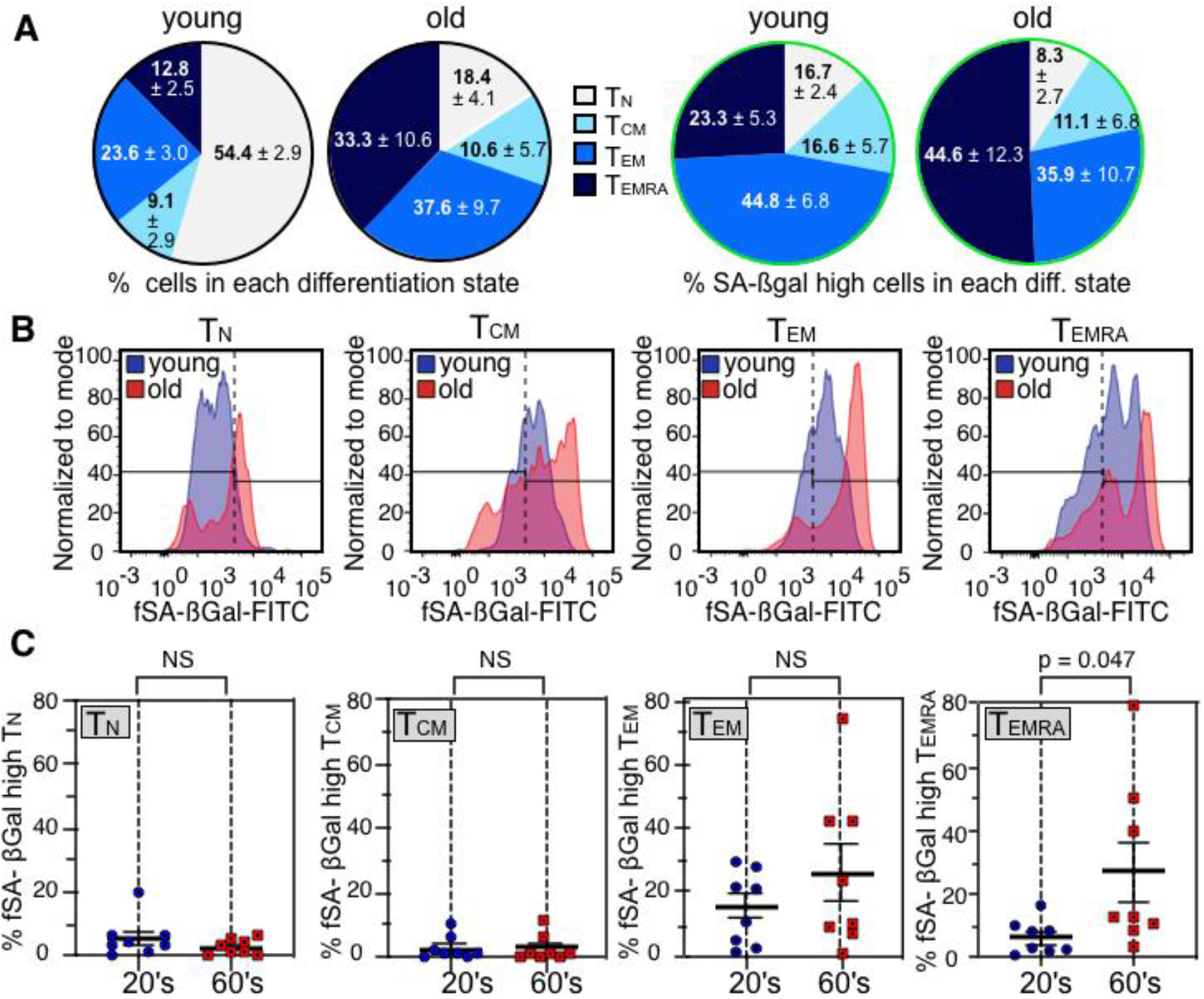
Senescent CD8+ T cells develop primarily in TEM and TEMRA subsets, (**A**) Distribution of indicated T cell differentiation states from young (left; n = 8) and old (right; n = 8) donors in unsorted (left) and fSA-ßGal high-sorted CD8+ T cell populations (right). (**B**) Representative fSA-ßGal intensity profiles and gates used to quantify fSA-ßGal high cells for indicated CD8+ T cell differentiation states from a young (blue) and old (red) donor. (**C**) Quantification of the percentages of fSA-ßGal high cells in young (and old donors for indicated CD8+ T cell differentiation states. Statistical significance was determined by an unpaired, two-tailed Student’s t test. NS: not significant.

### Senescent CD8+ T cells are distinct from exhausted PD1+, CD57+, KLRG1+, and from T_EMRA_ cells

T cell exhaustion is a phenotype caused by repeated antigenic stimulation by pathogens, resulting in functional impairment and the inability to produce cytokines such as IL-2, TNFα, and IFNγ (*18*). Similar to T_EMRA_ cells, exhausted CD8+ T cells display characteristics of SCs such as shortened telomeres, absence of telomerase activity, and reduced proliferative capacity (*30*). In addition, markers such as KLRG1 and CD57 have been shown to be expressed on exhausted or senescent CD8+ T cells, yet whether any of these markers accurately labels senescent CD8+ T cells with high SA-ßGal activity remains unclear(*18, 20*). To resolve this issue, we conducted t-distributed Stochastic Neighbor Embedding (t-SNE) analysis on CD8+ T cells from young and old donors that were simultaneously treated with the fSA-ßGal substrate and immunostained with various T cell exhaustion markers. This analysis revealed that only a small fraction of CD8+ T cells with high SA-ßGal activity expressed the inhibitory receptor PD1, a key feature of exhausted T cells (Fig. 5A). In fact, PD1 was expressed to a large degree also on CD8+ T cell with low fSA-ßGal signal intensities, demonstrating that T cell exhaustion is not equivalent to T cell senescence. Similarly, none of the other markers that have previously been reported to label CD8+ T cells with features of senescence, including CD57, KLRG1, or CCR7-/CD45RA+ (TEMRA), were expressed exclusively on CD8+ T cells with high fSA-ßGal signal intensities, nor did they label all cells that displayed high levels of fSA-ßGal signal intensities (Fig. 5A). Furthermore, TEMRA cells, generally regarded as a T cell differentiation stage that most closely resembles a state of cellular senescence, displayed a gene expression signature that was markedly distinct to that from senescent CD8+ T cells with high fSA-ßGal activity, as merely 53 % of DEGs were shared between these two cell states (Fig. 5B-C and Fig. S6). In fact, senescent CD8+ T cells have a transcriptome that is substantially different from any of the known CD8+ T cell differentiation states (Fig. S6), supporting our conclusion that cellular senescence of human CD8+ T cells in peripheral blood represents a unique T cell fate.

**Fig. 5.**
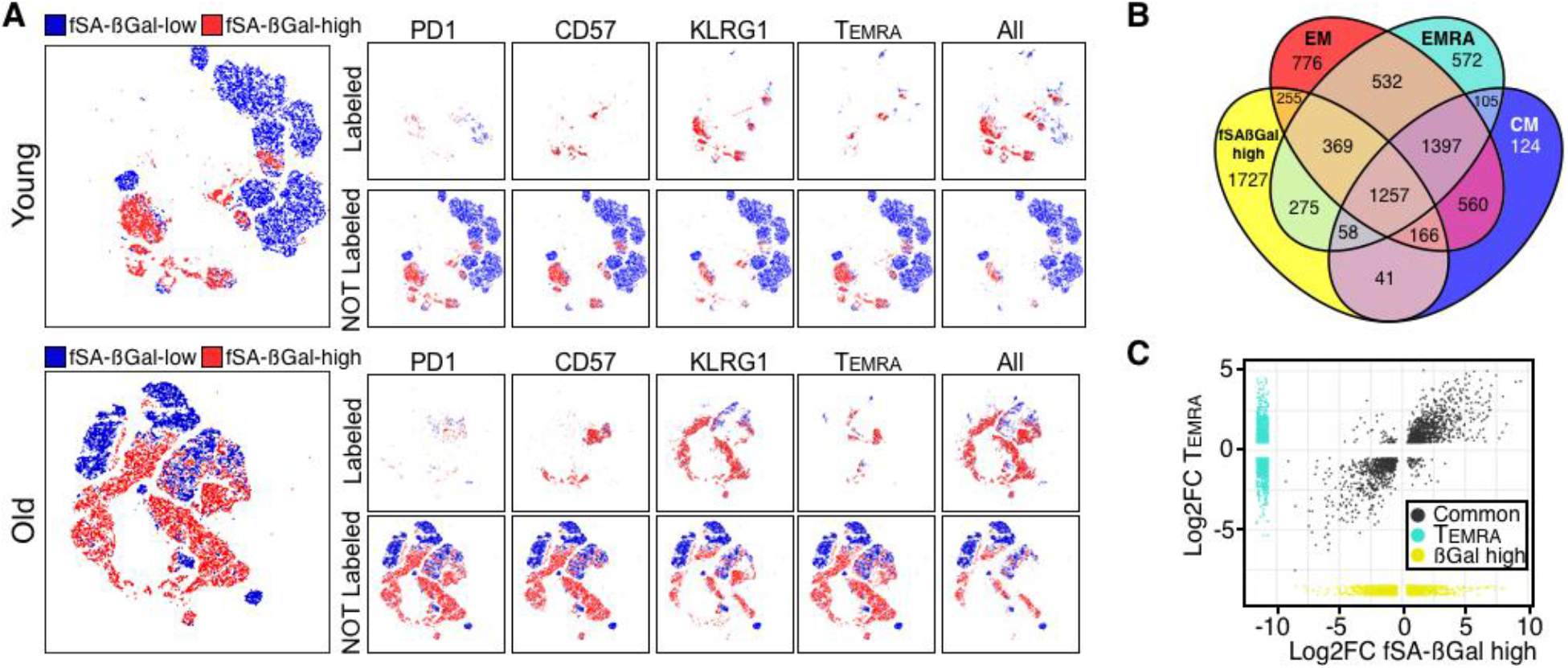
CD8+ T cells with high fSA-ßGal signal intensities are phenotypically distinct from exhausted and senescence-like cells and transcriptionally distinct from TEMRA cells. (**A**) t-SNE projections of CD8+ T cells with low (blue) and high (red) fSA-ßGal signal intensities that were also labeled with indicated cell surface receptors. Grey: CD8+ T cells that immunostained for indicated cell surface antigens but that did not fall into the fSA-ßGal-low or -high populations. (**B**) Venn diagrams portraying the intersections and disjunctive unions of DEGs in fSA-ßGal-high, T_CM_, T_EM_, and T_EMRA_ CD8+ T cells. (**C**) Correlation plot of the log2 fold changes of the DEGs in fSA-ßGal-high and EMRA CD8+ T cells. Dark grey points represent genes expressed in both populations. Turquoise dots are EMRA-specific genes. Yellow dots are fSA-ßGal-high-specific genes.

## DISCUSSION

Cellular senescence is a state in which a cell loses the ability to proliferate permanently, undergoes dynamic intracellular changes that stabilize the proliferative arrest, and develops pronounced secretory activity to signal and influence its surrounding environment (*1*). Given that CD8+ T cells, previously characterized as senescent based on expression of certain cell surface markers, still maintain some function and regain the ability to proliferate under certain conditions, it remains possible that T cell senescence, and potentially also senescence of other PBMC subsets, is reversible and therefore different compared to cellular senescence of other cell types. More likely, however, is that cell surface markers do not accurately distinguish senescent from non-senescent T cells *in vivo*. Our observations that CD8+ T cells *in vivo* develop defining features of cellular senescence, such as high levels of SA-ßGal activity, p16^INK4a^, macroH2A, dysfunctional telomeres, and a transcriptional signature that overlaps 71% with that of human fibroblasts that underwent replicative senescence *in vitro* for extended periods, support our conclusion that a true, irreversible and classical senescence response of CD8+ T cells exists. While we cannot exclude the possibility that CD8+ T cells with these senescence features retain the ability to proliferate under certain conditions, this scenario seems unlikely as a minimum of two senescence pathways, the p53/p21 and the p16^INK4a^/pRb pathway, would have to be inactivated for cells to re-enter the cell cycle. We therefore conclude that high levels of SA-ßGal activity is an accurate and reliable marker for cellular senescence of CD8+ T cells and other PBMCs.

The decline of immune function with age has been attributed to a number of factors, including to the age-associated impairment in thymopoiesis, a reduction in the ratio of naïve to memory cells, and an increasing chronic low-grade inflammation. These changes are hallmarks of “immunosenescence” and are considered to be the primary cause of the increased frequency and severity of disease and infections in the elderly. Surprisingly, impaired immune function can also accelerate the aging process, as it promotes the accumulation of senescent cells in a mouse model with defects in NK and T cell function, thereby causing animals to develop age-associated disorders and shortened lifespan(*31*). Further supporting the conclusion that impaired T cell function promotes aging in mammals is a recent study demonstrating that mice with defective CD8+/CD4+ T cells, due to T cell specific knockout of the mitochondrial transcription factor A (Tfam), accumulated senescent cells in various tissues in an accelerated manner, which caused these mice to develop numerous age-associated disorders and dramatically shortened their lifespan (*32*). Significantly, Tfam-deficient T cells displayed key features of cellular senescence, such as increased lysosomal content, reduced lysosomal pH, deficiency in cell proliferation, a reduced NAD+/NADH ratio, and increased production of pro-inflammatory cytokines TNFα and IFNγ, among others (*33*). The systemic senescence-inducing and pro-aging effects of these defective T cells were partially caused by their secreted molecules, as blocking TNFα suppressed development of senescent cells in various tissues and rescued many of the age-associated disorders (*32*). Thus, maintaining functional T cells throughout life appears to be critically important, not only because T cells target and eliminate senescent cells that increasingly develop in aging tissues (*31, 34*), but also because defects in T cell function that lead them to develop senescence-like features and increased production of pro-inflammatory cytokines such as TNFα, exacerbates the systemic senescent cell burden and promotes aging in mammals (*32*).

Remarkably, most of the reported characteristics of Tfam-deficient mouse T cells, including substantially increased expression of inflammatory cytokines TNFα and IFNγ are shared also with senescent human CD8+ T cells, as our study reveals. The observations that on average 64% of CD8+ T cells are senescent in healthy subjects in their 60s, therefore suggest that T cell senescence is a major contributing factor to immunosenescence with potentially profound physiological consequences for humans. We propose that the large abundance of senescent CD8+ T cells in peripheral blood of aged humans not only contributes to the increased susceptibility of the elderly to infections, but it likely also impairs clearance of senescent cells that accumulate in various tissues with age, accelerates development of systemic senescence through the active secretion of inflammatory SASP components and thereby promotes the development of age-associated pathologies and aging.

Our observations that telomeres are involved in senescence of CD8+ T cells opens opportunities to suppress immune cell senescence and improve immune surveillance in advanced age. This could be accomplished, for example, by enhancing telomerase activity or by reducing formation of dysfunctional telomeres using pharmacological means. Furthermore, neutralizing the activity of pro-inflammatory cytokines that are produced by senescent human CD8+ T cells such as TNFα, which has proven to be effective in suppressing systemic senescence caused by senescence-like T cells in mice (*32*), might also prove effective in suppressing the potentially damaging effects of senescent T cells in humans. Such strategies could potentially suppress age-associated diseases and enhance immune responses towards infections. In addition, the ability to efficiently detect, quantify, isolate, and analyze senescent PBMCs from human donor blood may prove useful for diagnostic purposes, to serve not only as a biomarker of biological age, but also as a potential marker of acute and chronic diseases.

## Acknowledgments

UH was supported by grants from the NCI (R01CA136533) and the NIA (R21AG067368) of the National Institutes of Health. PFB was supported by the NIAID (R01AI026806), (R01AI106), and the NIA (R21AG067368). The content is solely the responsibility of the authors and does not necessarily represent the official views of the National Institutes of Health. R.I.M-Z. is a Mexican National Scientific and Technology Council (CONACYT) and Mexican National Researchers System (SNI) fellow. HKD is supported by National Institutes of Health Integrated T32 Training Program in Infection, Immunity and Inflammation (Grant # 5T32AI125185);

## Author contributions

UH, PFB, RIM-Z and HD conceived the study. RIM-Z conducted experiments involving senescent human fibroblasts, RT-qPCR, and EdU incorporations and analyzed RNA-sequencing data. HKD collected blood, conducted FACS analysis and sorting of peripheral blood mononuclear cells and performed T cell proliferation assays. TV performed immunofluorescence analysis and quantification of senescence markers in sorted T cells. LGW provided cord blood and insight into neonatal physiology. UH, RIM-Z and HD assembled the figures and UH and RIM-Z wrote the manuscript with input from all authors.

## Competing interests

Authors declare no competing interests.

## Data and materials availability

All data is available in the main text or the supplementary materials.

## Supplementary Materials

Materials and Methods

Figures S1-S6

## Notes

### Competing Interest Statement

The authors have declared no competing interest.

## REFERENCES

1. A. Hernandez-Segura, J. Nehme, M. Demaria, Hallmarks of Cellular Senescence. Trends Cell Biol 28, 436–453 (2018).

2. M. Rhinn, B. Ritschka, W. M. Keyes, Cellular senescence in development, regeneration and disease. Development 146, (2019).

3. L. Prata, I. G. Ovsyannikova, T. Tchkonia, J. L. Kirkland, Senescent cell clearance by the immune system: Emerging therapeutic opportunities. Semin Immunol, (2019).

4. D. J. Baker et al., Naturally occurring p16(Ink4a)-positive cells shorten healthy lifespan. Nature 530, 184–189 (2016).

5. D. Munoz-Espin, M. Serrano, Cellular senescence: from physiology to pathology. Nat Rev Mol Cell Biol 15, 482–496 (2014).

6. J. J. Goronzy, C. M. Weyand, Mechanisms underlying T cell ageing. Nat Rev Immunol, (2019).

7. L. Pangrazzi, B. Weinberger, T cells, aging and senescence. Exp Gerontol 134, 110887 (2020).

8. M. Fumagalli et al., Telomeric DNA damage is irreparable and causes persistent DNA-damage-response activation. Nature cell biology 14, 355–365 (2012).

9. G. Hewitt et al., Telomeres are favoured targets of a persistent DNA damage response in ageing and stress-induced senescence. Nat Commun 3, 708 (2012).

10. I. Ben-Porath, R. A. Weinberg, The signals and pathways activating cellular senescence. The international journal of biochemistry & cell biology 37, 961–976 (2005).

11. U. Herbig, W. A. Jobling, B. P. Chen, D. J. Chen, J. M. Sedivy, Telomere Shortening Triggers Senescence of Human Cells through a Pathway Involving ATM, p53, and p21(CIP1), but Not p16(INK4a). Molecular cell 14, 501–513 (2004).

12. J. P. Coppe et al., Senescence-associated secretory phenotypes reveal cell-nonautonomous functions of oncogenic RAS and the p53 tumor suppressor. PLoS Biol 6, 2853–2868 (2008).

13. R. Zhang et al., Formation of MacroH2A-containing senescence-associated heterochromatin foci and senescence driven by ASF1a and HIRA. Dev Cell 8, 19–30 (2005).

14. R. Zhang, W. Chen, P. D. Adams, Molecular dissection of formation of senescence-associated heterochromatin foci. Molecular and cellular biology 27, 2343–2358 (2007).

15. D. J. Kurz, S. Decary, Y. Hong, J. D. Erusalimsky, Senescence-associated (beta)-galactosidase reflects an increase in lysosomal mass during replicative ageing of human endothelial cells. Journal of cell science 113 (Pt 20), 3613–3622 (2000).

16. V. Gorgoulis et al., Cellular Senescence: Defining a Path Forward. Cell 179, 813–827 (2019).

17. R. Vicente, A. L. Mausset-Bonnefont, C. Jorgensen, P. Louis-Plence, J. M. Brondello, Cellular senescence impact on immune cell fate and function. Aging Cell 15, 400–406 (2016).

18. J. P. Chou, R. B. Effros, T cell replicative senescence in human aging. Curr Pharm Des 19, 1680–1698 (2013).

19. Y. Song et al., T-cell Immunoglobulin and ITIM Domain Contributes to CD8(+) T-cell Immunosenescence. Aging Cell 17, (2018).

20. W. Xu, A. Larbi, Markers of T Cell Senescence in Humans. Int J Mol Sci 18, (2017).

21. G. Dimri et al., A biomarker that identifies senescent human cells in culture and in aging skin *in vivo*. Proc. Natl. Acad. Sci. USA 92, 9363–9367 (1995).

22. F. Debacq-Chainiaux, J. D. Erusalimsky, J. Campisi, O. Toussaint, Protocols to detect senescence-associated beta-galactosidase (SA-betagal) activity, a biomarker of senescent cells in culture and in vivo. Nat Protoc 4, 1798–1806 (2009).

23. P. Langfelder, S. Horvath, WGCNA: an R package for weighted correlation network analysis. BMC Bioinformatics 9, 559 (2008).

24. M. De Cecco et al., L1 drives IFN in senescent cells and promotes age-associated inflammation. Nature 566, 73–78 (2019).

25. R. I. Martinez-Zamudio et al., AP-1 imprints a reversible transcriptional programme of senescent cells. Nature cell biology, (2020).

26. S. H. Ross, D. A. Cantrell, Signaling and Function of Interleukin-2 in T Lymphocytes. Annu Rev Immunol 36, 411–433 (2018).

27. V. Appay, R. A. van Lier, F. Sallusto, M. Roederer, Phenotype and function of human T lymphocyte subsets: consensus and issues. Cytometry A 73, 975–983 (2008).

28. S. M. Henson et al., p38 signaling inhibits mTORC1-independent autophagy in senescent human CD8(+) T cells. J Clin Invest 124, 4004–4016 (2014).

29. L. A. Callender et al., Human CD8(+) EMRA T cells display a senescence-associated secretory phenotype regulated by p38 MAPK. Aging Cell 17, (2018).

30. M. Bellon, C. Nicot, Telomere Dynamics in Immune Senescence and Exhaustion Triggered by Chronic Viral Infection. Viruses 9, (2017).

31. Y. Ovadya et al., Impaired immune surveillance accelerates accumulation of senescent cells and aging. Nat Commun 9, 5435 (2018).

32. G. Desdin-Mico et al., T cells with dysfunctional mitochondria induce multimorbidity and premature senescence. Science, (2020).

33. F. Baixauli et al., Mitochondrial Respiration Controls Lysosomal Function during Inflammatory T Cell Responses. Cell Metab 22, 485–498 (2015).

34. B. I. Pereira et al., Senescent cells evade immune clearance via HLA-E-mediated NK and CD8(+) T cell inhibition. Nat Commun 10, 2387 (2019).

35. P. K. Puvvula et al., Long noncoding RNA PANDA and scaffold-attachment-factor SAFA control senescence entry and exit. Nat Commun 5, 5323 (2014).

36. M. Tsang, J. Gantchev, F. M. Ghazawi, I. V. Litvinov, Protocol for adhesion and immunostaining of lymphocytes and other non-adherent cells in culture. Biotechniques 63, 230–233 (2017).

37. P. L. Patel, U. Herbig, Detection of Dysfunctional Telomeres in Oncogene-Induced Senescence. Methods Mol Biol 1534, 69–78 (2017).

38. M. E. Ritchie et al., limma powers differential expression analyses for RNA-sequencing and microarray studies. Nucleic acids research 43, e47 (2015).

39. M. I. Love, W. Huber, S. Anders, Moderated estimation of fold change and dispersion for RNA-seq data with DESeq2. Genome Biol 15, 550 (2014).

40. A. Subramanian et al., Gene set enrichment analysis: a knowledge-based approach for interpreting genome-wide expression profiles. Proceedings of the National Academy of Sciences of the United States of America 102, 15545–15550 (2005).

41. A. Liberzon et al., The Molecular Signatures Database (MSigDB) hallmark gene set collection. Cell Syst 1, 417–425 (2015).

