## Supplemental Information for "Conclusive Identification of Senescent T Cells Reveals Their Abundance in Aging Humans"

**This PDF file includes:**

Materials and Methods
Figs. S1 to S6

### Materials and Methods

**Cell culture.** GM21 fibroblasts were obtained from the Coriell Institute and cultured in Dulbecco's Modified Eagle's medium (DMEM) containing 10% FBS and 1X penicillin/streptomycin at 37˚C and 3% oxygen. GM21-ER:RAS were generated by retroviral transduction as previously described(*35*). OIS was induced by addition of 200 nM 4-hydroxytamoxifen (4-OHT) for 14 days, with media and 4-OHT being replenished every 3 days. For DNA damage/etoposide-induced senescence, GM21 fibroblasts were treated with a single dose of 20 µM etoposide followed by incubation for 16 days. For both senescence models, SA-ßGal activity was detected using a self-immobilizing fSA-βGal substrate. Briefly, control and senescent cells were collected by trypsinization, washed 2X in complete DMEM, and resuspended at a concentration of 1 x $10^6$ cells/ml. Cell suspensions (1ml total volume) were subsequently incubated with 1X bafilomycin A1 solution for 1 hour at 37º C followed by incubation with the fSA-βGal substrate for an additional 30 min. Cells were centrifuged at 2,000 rpm for 4 min at RT, washed twice in PBS, and collected by an additional centrifugation step as described and analysed as described in the main text.

**EdU incorporation.** Proliferating and senescent GM21-ER:RAS fibroblasts as well as FACS-sorted fSA-ßGal-low and -high fibroblasts from mixed populations were seeded on collagen coated coverslips (Corning, Corning, NY) in 6-well plates, supplemented with DMEM 10% FBS and antibiotics (1X penicillin/streptomycin). EdU incorporation was performed using the Click-iT EdU Alexa Fluor Imaging Kit as per the manufacturer's instructions (Thermo Fisher Scientific, Waltham, MA). Cells were imaged using Images were acquired with a Zeiss Axio Observer Z1 Inverted Phase Contrast Fluorescence Microscope using a 10x objective and images analyzed using the with Zeiss ZEN 2.5 (blue edition) software.

**Peripheral Blood Mononuclear Cell Isolation.** The Institutional Review Board of New Jersey Medical School approved this study (ID Pro0119980237, approved November 14, 2019). Blood was obtained from healthy, consenting donors 22-65 years old in heparinized collection tubes. Peripheral blood mononuclear cells were separated using Lymphocyte Separation Medium (Corning, Manassas, VA) via density centrifugation following the manufacturer's protocol. PBMCs were cultured in RPMI 1640 (VWR Life Science, Radnor, PA) with L-glutamine (Corning, Manassas, VA) supplemented with 10% heat-inactivated fetal bovine serum (Gibco, Gaithersburg, MD), 100 U/mL penicillin, 100 mg/mL streptomycin, 100 mg/mL gentamicin, and 5 mM HEPES (Sigma Aldrich, St. Louis, MO). Cells were counted using the Cellometer Auto 2000 (Nexcelom, Lawrence, MA) and cultured at 2 x $10^6$ cells/ml.

**Antibodies.** APC anti-CD54RA (HI100), PE Cy7 anti-CCR7 (G043H7), PerCP Cy5.5 anti-CD8(SK1), PerCP Cy5.5 anti-CD8 (SK1), AF700 anti-CD3 (HIT3A), APC anti-CD3 (HIT3A), PE Cy7 anti-CD4 (OKT4), PerCP Cy5.5 anti-CD303 (210A), APC Cy7 anti-CD14 (HDC14), BUV395 anti-CD45 (H130), Brilliant Violet 510 anti-CD57 (QA17A04), Brilliant Violet 711 anti-PD1 (EH12.2H7), PE anti-KLRG1 (14C2A07), Pacific Blue anti-CD56 (5.1H11), AF700 anti-CD19 (HIB19) were purchased from BioLegend (San Diego, CA), anti-p16 (JC8, GeneTex, San Antonio, TX), anti-53BP1 (polyclonal; Novus, Littleton, CO), anti-macroH2A (H-39, Santa Cruz, Dallas, TX). Secondary antibodies used for immunofluorescence staining were : Cy3 donkey anti-mouse (Jackson ImmunoResearch, West Grove, PE) and AlexaFluor 488 goat anti-rabbit (Invitrogen, Waltham, MA).

**Surface staining**. PBMCs were incubated with a 1X solution of bafilomycin A1 before the addition of the fSA-βGal substrate. PBMCs were washed with PBS (Corning, Manassas, VA) and incubated with 5 µL of heat-inactivated purified human serum, collected from healthy donors, at room temperature for 5 minutes. Cells were then incubated for 20 minutes with antibody at 4 degrees and washed with 2% FCS (Gibco, Gaithersburg, MD), in PBS. To test for viability before cell sorting, cells were resuspended in 25 µL/mL 4',6-diamidino-2-phenylindole (DAPI (BioLegend, San Diego, CA) prior to sample collection. Samples for flow cytometric analysis were acquired at 300,000 events using an BD LSRFortessa X-20 (BD Biosciences, Franklin Lakes, NJ) and analysis was performed with FlowJo software (FlowJo, (versions 9.3.2, 10) LLC Ashland, OR).

**CD8+ T Cell Enrichment**. CD8+ T cells were from PBMCs prepared as described, by negative selection using the StemCell EasySep Human CD8+ T Cell Isolation Kit (Cambridge, MA) according to the manufacturer's protocol. Purity of enriched cells was assessed via flow cytometry as the proportion of CD3+ and CD8+ cells. Purities ranged from 95-98%. CD8+ T cells were counted using the Cellometer Auto 2000 (Nexcelom, Lawrence, MA) and cultured at $2 \times 10^6$ cells/ml. Cells were sorted based on fSA-ßGal intensity on the BD FACSAria Fusion (BD Biosciences, Franklin Lakes, NJ).

**CD8+ T Cell Proliferation.** 50,000 sorted CD8+ T cells were stained with CellTrace Violet (ThermoFisher, Waltham, MA) following manufacturer's protocol. Labeled cells were mixed with 400,000 unlabeled autologous PBMCs and cultured in 200 µl media with 2 µg/ml plate-bound anti-CD3 (OKT3) and 5 µg /ml soluble anti-CD28 (CD28.2) purchased from Biolegend (San Diego, CA). After 5 days, cells were collected and stained with Zombie UV Fixable Viability Kit (Biolegend, San Diego, CA), anti-CD3 and anti-CD8. Samples were acquired on a X-20 BD LSRFortessa (BD Biosciences, Franklin Lakes, NJ). Sorted cells were identified as all CD3+ CD8+ CellTrace Violet+ cells.

**Senescence gene expression profiling.** RNA from GM21 and CD8+ T cell populations described in this study was purified using a Macherey-Nagel RNA XS Plus Kit according to the manufacturer's instructions (Macherey-Nagel, Duren, Germany). cDNA was generated with a Biorad iScript cDNA synthesis kit according to the manufacturer's instructions (Biorad, Hercules, CA). For all genes assessed, Qiagen Quantitect primer assays were used and real-time RT-qPCR was performed on a Biorad CFX96 Real-Time PCR detection system (Biorad, Hercules, CA). Data were analyzed on the Biorad CFX Manager software 3.1.

**Statistical analysis.** With the exception of microarray and RNA-seq analyses, all statistical tests utilized in this study were performed in Graphpad Prism 8 and are indicated in the legend of their respective figure.

**Preparation of T cells for immunofluorescence staining.** Immunofluorescence staining of T cells was performed as described(*36*). Briefly, sorted CD8+ were collected and re-suspended in PBS at $10^6$ cells/ml. The cell suspension was layered on glass coverslips and was left standing for 30 minutes to let the cells attach on the coverslip through gravity sedimentation. After 30 minutes the cells were fixed in 4% formaldehyde for 15 minutes and permeabilized for 15 minutes with 0.2% Triton-X PBS (PBST 0.2%). Following permeabilization, cells were incubated in blocking buffer (4% BSA in PBST 0.1%) for 1 hour.

**Immunofluorescence staining (IF).** Cells were incubated with primary antibodies diluted in blocking buffer for 2 hours at room temperature and were subsequently washed 3 times (7 minutes each) with PBST 0.1%. Incubation with secondary antibodies was performed for 1 hour at room temperature followed by 3 washes (7 minutes each) with PBS. Cells were then air dried and mounted on slides using DAPI containing mounting medium (Invitrogen, Waltham, MA).

**Immunofluorescence staining (IF) combined with telomere FISH.** Telomere-ImmunoFISH was performed as described in (*37*). Briefly, following the blocking step, cells were washed 3 times with PBS (5 minutes each) and were dehydrated by subsequent submersion in 70%, 90% and 100% ethanol (3 minutes each). After dehydration cells were left to complete air dry and were incubated with 0.5 µg/ml TelC-Cy3 PNA probe (Panagene, Korea) in hybridization buffer (70% formamide, 12 mM Tris-HCl pH=8.0, 5 mM KCl, 1mM MgCl2, 0.001% Triton X-100, 0.25% acetylated BSA) for 5 minutes at 80 $^{o}$C, to allow denaturation of DNA and subsequent hybridization of the telomere specific PNA probe. The cells were then left to cool gradually and further incubated overnight in a humidified chamber at room temperature. After hybridization cells were washed 2 times with 70% formamide/0.6X SSC (15 minutes each) followed by 2 washes with 2X SSC buffer (10 minutes each). After the last wash immunostaining was performed as described above using anti- 53BP1 primary antibodies.

**Image acquisition and analysis of DNA damage and Telomere-dysfunction Induced DNA damage Foci (TIF).** Images were acquired with a Zeiss Axio Observer Z1 Inverted Phase Contrast Fluorescence Microscope using a 63X oil-immersive lens. Images were obtained using Z-stacks and were analyzed with Zeiss ZEN 2.5 (blue edition) software. For each experiment at least 100 cells were analyzed and the number of 53BP1 foci per cell, as well as the co-localization of 53BP1 foci with telomeres, were assessed. Only distinct, well-formed 53BP1 foci in each plain focus of the Z-stack were counted for absolute number of foci per cell, as well as for colocalizations. T cells with only granular non-focal staining pattern of 53BP1 were excluded from analysis.

**RNA-seq.** RNA from sorted CD8+ fSA-ßGal-low and -high from young and old donors (3 each) was purified using a Macherey-Nagel RNA XS Plus kit according to the manufacturer's instructions (Macherey-Nagel, Duren, Germany). RNA integrity was evaluated in a Bioanalyzer 2100 system and only RNA with an integrity of number of >= 7 was used for library preparation. Libraries were constructed using the SMARTer Stranded V2 (TakaraBio) according to the manufacturer's instructions (TakaraBio, Mountain View, CA). Paired-end sequencing was performed on an Illumina HiSeq 2500 instrument. At least 40 million reads per sample (20 million per strand) were obtained and used for downstream analyses.

**Analysis of microarray data from Callender et al., 2018(*29*).** Publicly available Affymetrix HuGene 2.0 microarrays for N, CM, EM and EMRA CD8+ T cells (6 each)(*29*) cells were analyzed using open-source Bioconductor packages on R. All microarrays were normalized simultaneously using the robust multi-array normalization algorithm implemented in the *oligo* package. After removing internal control probes, deciles of average expression were independently defined for each CD8+ T cell population. Probes falling in the lowest 4 deciles of expression were removed. Identification and removal of unidentified sources of variability were performed using the *sva* package. Probes were annotated by combining the *HuGene 2.0* and *bioMart* annotations. Hierarchical clustering and principal component analysis were used to monitor the behavior of the datasets at every step of the pre-processing. Differential expression was performed using *limma* with the default parameters, comparing the overall expression changes of CM, EM and EMRA relative to N CD8+ T cells. The statistical approach involves empirical Bayes-moderated *t*-statistics applied to all contrasts for each probe, followed by moderated *F*-statistic to tests whether all the contrasts are zero, evaluating the significance of the expression changes observed(*38*). For significant probes, *p*-values were corrected for multiple testing with the FDR approach using a significance level of 0.005 as a cut-off. Probes matching these criteria that also exhibited a minimal absolute fold change of 1.4 were considered for downstream analyses.

**RNA-seq analysis.** FASTQ files were aligned to the version 19 of the human reference genome (hg19) using *bowtie2* using the local alignment option (bowtie2 --local -x ./hg19/hg19) which performs soft clipping of read characters to maximize alignment scores. Alignments were further processed using *samtools v1.9.* For genome track visualizations, alignments were normalized using *deeptools v3.3.1* using the RPGC approach to obtain 1X coverage (bamCoverage -b --normalizeUsing RPGC --effectiveGenomeSize 2864785220 --ignoreDuplicates --binSize 10 --verbose -o). Reads per exon (hg19) were quantified using the *summarizeOverlaps* package and genes with at least 10 reads in at least 3 samples were kept. Correction of batch effects was performed with *sva*. Data transformation (regularized-log [rld] transformation), exploratory visualization (PCA and hierarchical clustering) and differential expression analysis were performed with *DESeq2* using the default parameters as previously described(*39*) and base R functions. Differentially expressed genes identified by this approach (absolute fold change of 1.25) were considered for downstream analyses.

**Transcriptional co-expression networks and functional overrepresentation.** For microarray and RNA-seq datasets analyzed, the debatched normalized fluorescence and rld-transformed counts, respectively, of the DEGs identified were used as input for the *WGCNA(23)* package and clustered using the 'signed' option with default parameters with the exception of the soft-thresholding power, which was set to 18 for microarray and 19 for RNA-seq DEGs, respectively. The co-expression modules identified for each dataset were visualized with heatmaps, correlation and network plots using *WGCNA* and *pheatmap*, and functionally characterized by overrepresentation tests with the *clusterProfiler* package using the Molecular Signature Database Hallmark gene sets(*40, 41*). For the comparative analysis of microarray and RNA-seq datasets of CD8+ T cells RNA-seq datasets from this study with datasets from De Cecco et al.(*24*) and Callender et al. (*29*), the lists of DEGs derived from the *WGCNA* modules from the microarray and RNA-seq datasets were initially characterized for intersections and disjunctive unions using *eulerr*, allowing for the identification of cell state- and cell type-specific DEGs. These lists were functionally characterized with *clusterProfiler* as described above. For a quantitative visualization of the intersections and disjunctive unions of DEGs of each CD8+ T cell state from Callender et al., we displayed the pair-wise correlation of the log2 fold change of DEGs common to CD8+ fSA-ßGal-high and CD8+ T cells at the CM, EM and EMRA stages, respectively. R scripts are available upon request.

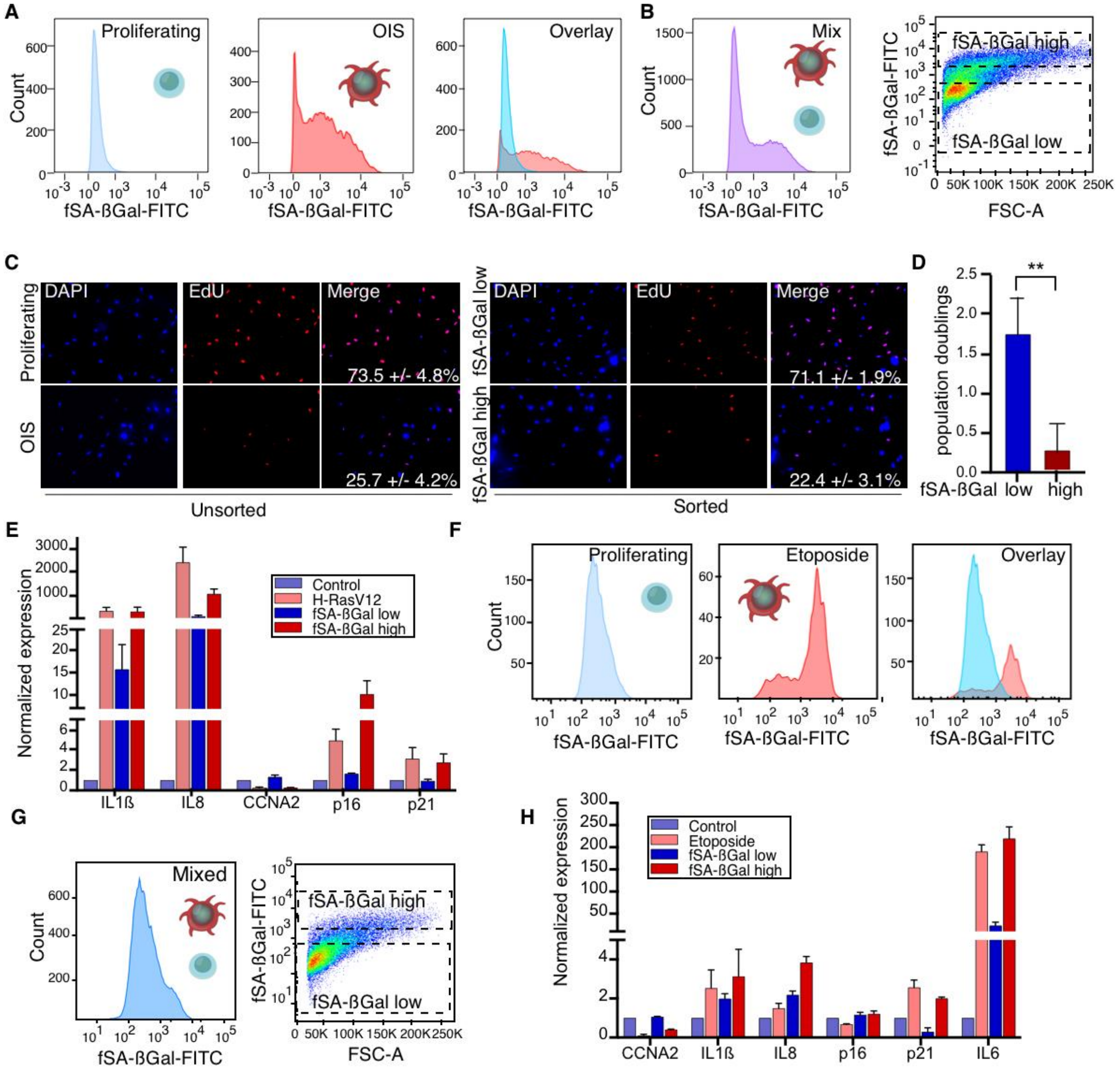


**Fig. S1.**
Senescent human fibroblasts can be accurately isolated from mixed cell populations using a fluorogenic SA-βGal substrate. (**A**) Representative flow cytometry fSA-ßGal intensity profiles for proliferating and oncogene-induced senescent (OIS) fibroblasts. Overlay of the two histograms is shown to the right. (**B**) Proliferating fibroblasts and fibroblasts in OIS were mixed at a ratio of 3:1 and subjected to the fSA-βGal staining procedure. Histogram: fSA-ßGal intensity profile for mixed cell population; dot-plot: gating strategy to sort for fibroblasts with low and high fSA-ßGal signal intensities as indicated. (**C**) Immunofluorescence analysis of EdU incorporation (red) of non-sorted proliferating and OIS fibroblasts (left) and fibroblasts sorted by FACS based on gates shown in b (right), as indicated. Percentages of EdU positive cells are indicated (average and S.E.M of 2 independent experiments are shown). (**D**) Quantification of population doublings of fibroblasts sorted by FACS based on gates shown in b 4 days after re-plating. n = 3 +/-S.E.M. Statistical significance was calculated with an unpaired, two-tailed t-test. ** p = 0.0124 (**E**) Representative RT-qPCR expression profiles of indicated senescence-associated genes were determined for proliferating (control), OIS (H-RasV12), sorted, as in b, fSA-ßGal-low and fSA-ßGal-high fibroblasts. n = 3. (**F**) as (a) using etoposide-induced senescent fibroblasts. (**G**) as (b) using etoposide-induced senescent

fibroblasts. **h**, RT-qPCR expression profiles of indicated senescence-associated genes as in (e). n = 2.

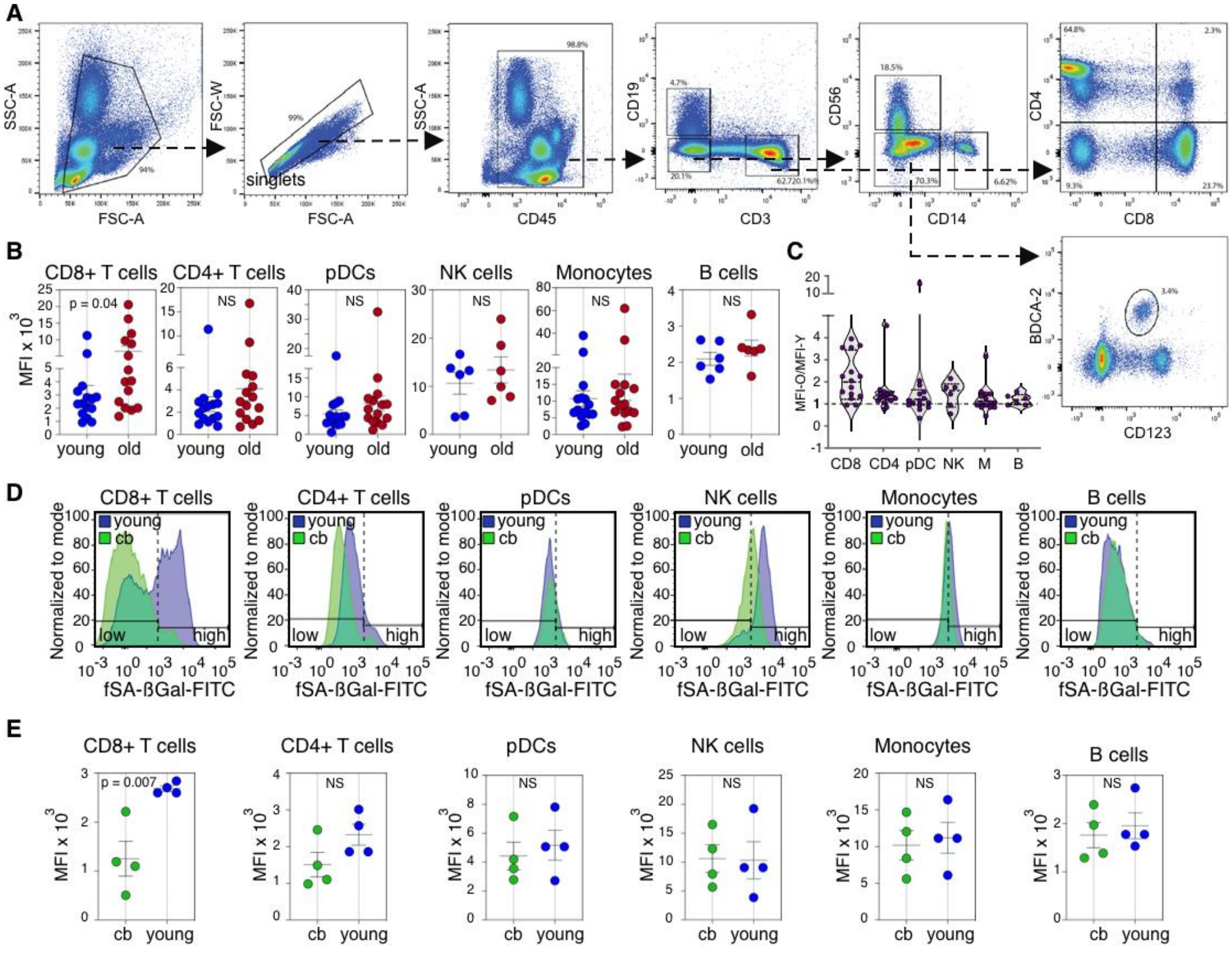

**Fig. S2.**

fSA-βGal fluorescence signal intensities of human T lymphocytes show consistent increases with age. (**A**) Gating strategy to identify PBMCs, single cell events, CD45+, CD56+, CD14+, CD19+ CD4+ and CD8+ T cells, and CD123+ BDCA2+ cells. (**B**) Mean fluorescent fSA-βGal signal intensities (MFI; mean +/- S.E.M) of indicated PBMC subsets from young and old donors. (**C**) Violin plot showing ratios of MFIs between old and young donor pairs in indicated PBMC subsets. (**D**) Representative fSA-ßGal intensity profiles and gates used to quantify fSA-ßGal high cells for indicated PBMC subsets from cord blood (green) and young (blue) donors. (**E**) Mean fluorescent fSA-βGal signal intensities (MFI) of indicated PBMC subsets from analyzed cord blood and young donors as indicated.

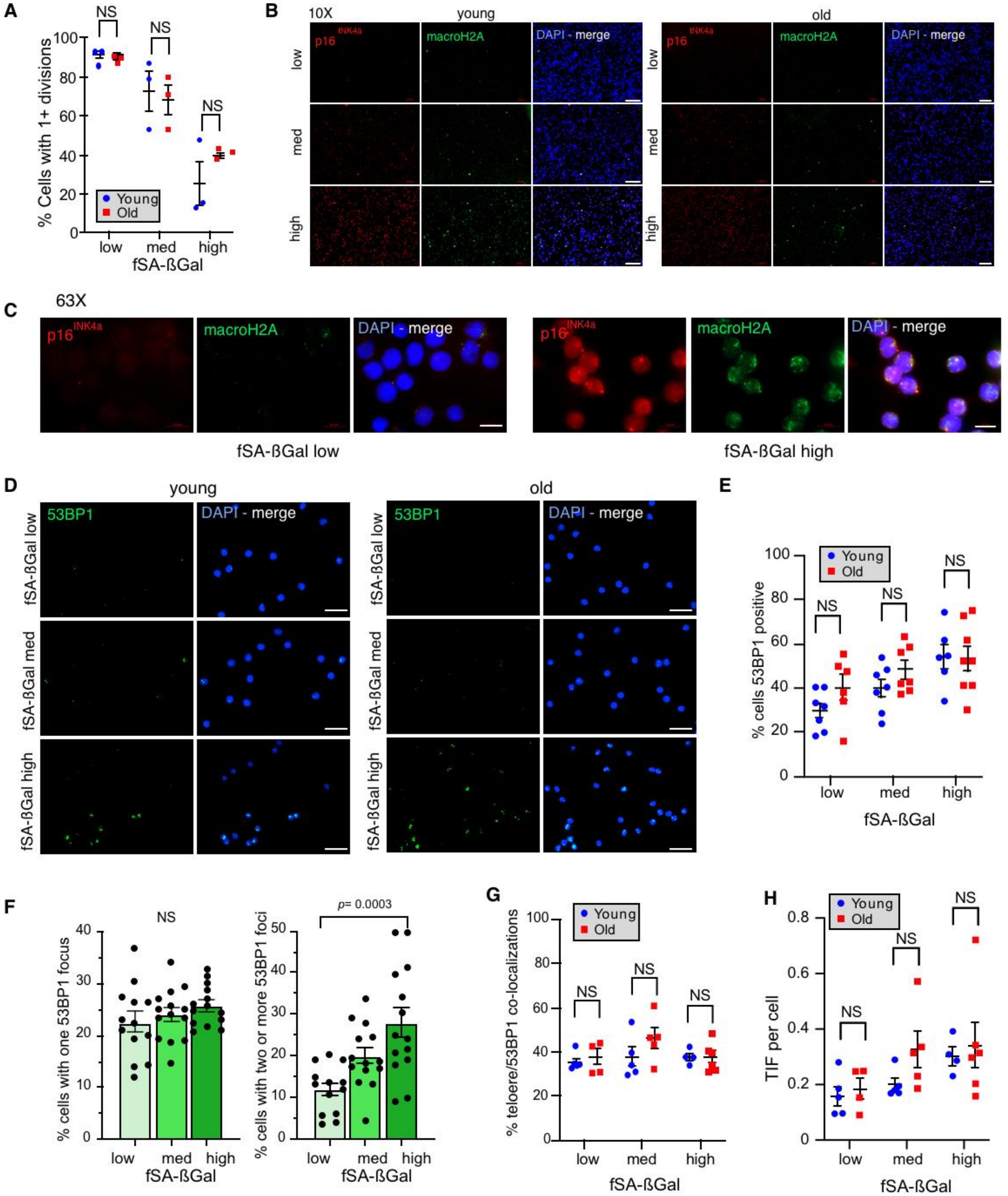


**Fig. S3.**

CD8+ T cells with high fSA-βGal signal intensities display hallmark features of senescent cells. (**A**) Quantification of more than one cell divisions of CD8+ T cells from 3 young (blue) and 3 old (red) donors following anti-CD3 and anti-CD28 stimulation for 60 h. Statistical significance was calculated with a two-way ANOVA. NS: not significant. (**B**) Immunofluorescence analysis of CD8+ T cells from a young and old donor sorted based on fSA-βGal signal intensities, as indicated, using antibodies against macroH2A (green) and p16$^{INK4a}$ (red). Blue: DAPI. Images were acquired at 10X magnification (scale bar 100 μm) and (**C**) at 63X magnification (scale bar 10 μm) (**D**) Immunofluorescence analysis as in B using antibodies against 53BP1 (green). Blue: DAPI. (scale bar 20 μm) (**E**) Quantification of CD8+ T cells positive for 53BP1 foci, sorted based on indicated fSA-βGal signal intensities from young (blue) and old (red) donors. Statistical significance was calculated with a two-way ANOVA. NS: not significant. (**F**) Quantification of CD8+ T cells sorted based on indicated fSA-βGal signal intensities from young and old donors combined, displaying one 53BP1 focus only (left bar graph) and two or more 53BP1 foci (right graph). Young n = 8, Old n=8. Bars and whiskers depict mean +/- S.E.M. Statistical significance was calculated with a one-way ANOVA. NS: not significant. (**G**) Quantification of percent 53BP1-telomere colocalizations as indicated. (**H**) Quantification of TIF per cell in CD8+ T cells sorted based on indicated fSA-βGal signal intensities from young (blue) and old (red) donors. Statistical significance was calculated with a two-way ANOVA (g,h). NS: not significant.

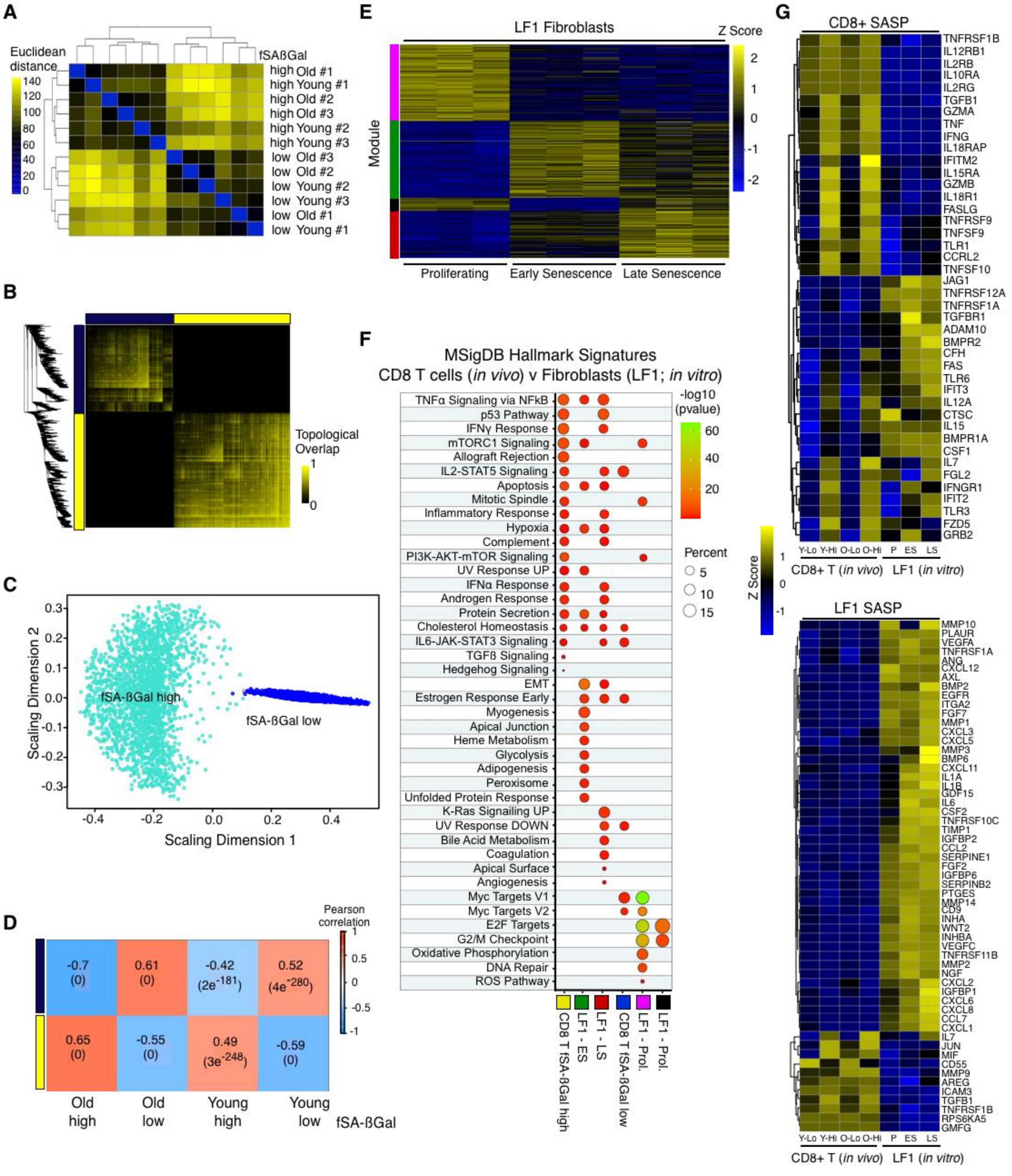


**Fig. S4.**

Gene expression profiles of senescent human CD8+ T cells and senescent human lung fibroblasts (LF1) (**A**) Hierarchical clustering of r-log-regularized counts per exon (see Methods) of CD8+ fSA-ßGal-low and -high T cells from young and old donors. (**B**) Network plot of the 4,149 DEGs in CD8+ fSA-ßGal-high T cells. Note that two distinct modules are identified (dark blue and yellow). (**C**) Multidimensional scaling plot of the two gene modules (fSA-ßGal-low and -

high) identified in Fig. 3b. Each dot represents a single gene. (**D**) Correlation plot showing gene module usage in CD8+ fSA-ßGal-low and -high CD8+ T cells of young and old donors as indicated. (**E**) Heatmap showing four modules of co-expressed genes specific for the indicated human LF1 fibroblast samples using an unsupervised WGCNA clustering approach. (**F**) Functional overrepresentation analysis comparing enrichment of MSigDB hallmark gene sets in CD8+ fSA-ßGal-low and -high T cells, as well as proliferating, early replicative senescent (ES; 4 months) and late replicative senescent (LS) human LF1 fibroblasts. Circles are color-coded according to the FDR-corrected p-value based on the hypergeometric distribution test. (**G**) Expression heatmaps of a selection of CD8+ T cell (top) and LF1- (bottom) specific SASP genes in low (lo) and high (hi) fSA-ßGal CD8+ T cells from young and old donors and LF1 fibroblasts in proliferation (P), early (E; 2 months) and late (L; 4 months) senescence. Each column represents the average expression of 3 independent donors (CD8+) and 3 independent experiments (LF1).

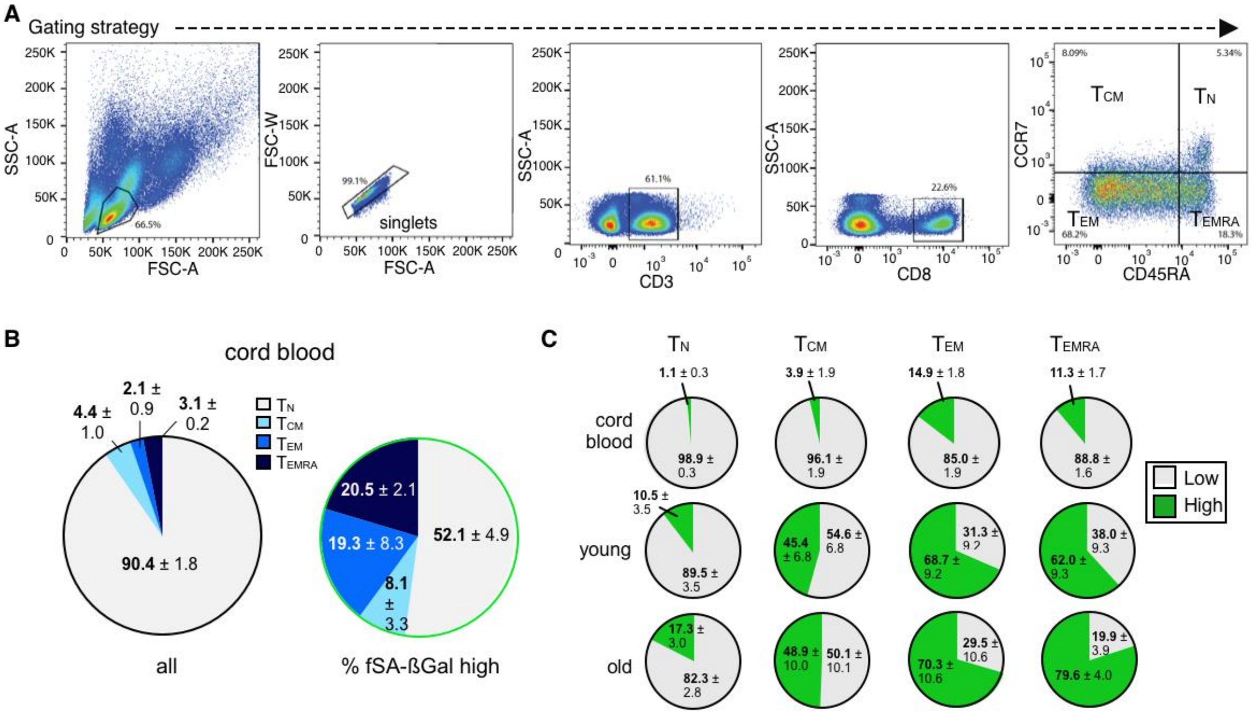


**Fig. S5.**

Senescent CD8+ T cells can be detected in cord blood and senescent CD8+ T cells are not exhausted. (**A**) Gating strategy to identify CD8+ T cell subsets by expression of CCR7 and CD45RA. (**B**) Distribution (% +/- S.E.M.) of indicated T cell differentiation states from cord blood (n = 3) in unsorted (top) and fSA-βGal high-sorted CD8+ T cell populations (bottom). (**C**) Fraction of fSA-βGal low (grey) and fSA-βGal high (green) cells within indicated CD8+ T cell differentiation states from cord blood (top row), young donors (middle row) and old donors (bottom row).

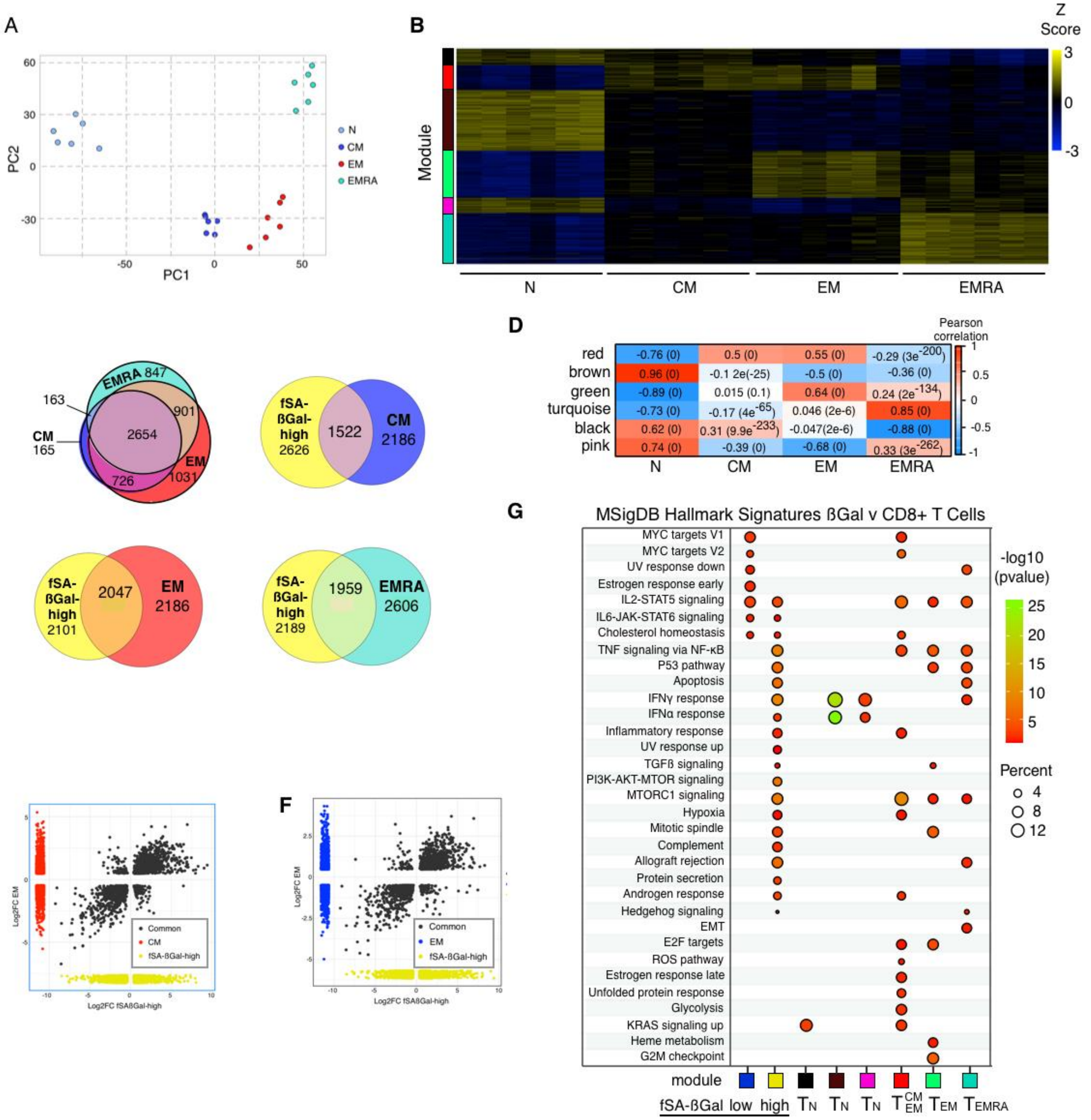


**Fig. S6.**

Senescent CD8+ T cells are transcriptionally distinct from TN, TCM, TEM, and TEMRA cells. (**A**) Principal component analysis of normalized and debatched transcriptomes of N, CM, EM and EMRA CD8+ T cells. CD8+T cell differentiation stages are color-coded. Microarray data from Callender et al, 2018., were reanalyzed (a-g). (**B**) Expression heatmap of the 5,593 DEGs across each CD8+ T cell differentiation stage. Microarray data from Callender et al, 2018 were clustered using WGCNA. Each column represents an individual donor. Data are represented as Z-scores. (**C**) Venn diagrams portraying the intersections and disjunctive unions of DEGs in fSA-ßGal-high $T_{CM}$, $T_{EM}$, and $T_{EMRA}$ CD8+ T cells. (**D**) Correlation plot showing gene module usage in $T_N$, $T_{CM}$, $T_{EM}$, and $T_{EMRA}$ CD8+ T cells. (**E, F**) Correlation plot of the log2 fold changes of the DEGs in fSA-ßGal-high, CM (e) and EM (f) CD8+ T cells. Dark grey points represent genes expressed in both populations for each pair-wise comparison. Red dots are CM-specific genes.

Blue dots are EM-specific genes. Yellow dots are fSA-ßGal-high-specific genes. (**G**) Functional overrepresentation analysis depicting enrichment of Molecular Signature Database gene sets in the fSA-ßGal-low and -high gene modules identified in Figure 3 and in differentiation stage-specific gene of CD8+ T cells. Note that CD8+ fSA-ßGal-high T cells enrich for most gene sets found in CD8+ T cells at every differentiation stage. Circles are color-coded according to the FDR-corrected p-value based on the hypergeometric distribution test.
